# Conserved units of co-expression in bacterial genomes: an evolutionary insight into gene regulation

**DOI:** 10.1101/012526

**Authors:** Ivan Junier, Olivier Rivoire

## Abstract

Genome-wide measurements of transcriptional activity in bacteria indicate that the transcription of successive genes is strongly correlated beyond the scale of operons. However, the underlying mechanisms are poorly characterized and a systematic method for identifying local groups of co-transcribed genes is lacking. Here, we identify supra-operonic segments of consecutive genes by comparing gene proximity in thousands of bacterial genomes. Structurally, the segments are contained within micro-domains delineated by known nucleoid-associated proteins, and they contain operons with specific relative orientations. Functionally, the operons within a same segment are highly co-transcribed, even in the absence of regulatory factors at their promoter regions. Hence, operons with no common regulatory factor can be co-regulated if they share a regulatory factor at the level of segments. To rationalize these findings, we put forward the hypothesis supported by RNA-seq data that facilitated co-transcription, the feedback of transcription into itself involving only DNA and RNA-polymerases, may represent both an evolutionary primitive and a functionally primary mode of gene regulation.

## Introduction

Differential gene expression underlies much of the dynamics of living cells. Describing globally gene coexpression, understanding its relation to genome organization, and explaining the mechanisms behind it are fundamental challenges in biology (Browning and Busby, 2004). These challenges have been met to a significant extent. Genome-wide measurements of transcriptional activity are available for different cells in different conditions, and many mechanisms are known which, in principle, can account for the observations (Browning and Busby, 2004). Yet, as reviewed below, the decomposition of bacterial genomes into operons and their regulation by sigma factors (SFs) and transcription factors (TFs), often presented as a first-order description of gene regulation in bacteria, fail to account for prominent features of gene expression. The presence of other regulatory factors is indeed well-recognized. These include small metabolites (Browning and Busby, 2004), small RNAs (Waters and Storz, 2009), transcriptional attenuators (Henkin and Yanofsky, 2002), global physiological effects (Klumpp *et al*, 2009; Berthoumieux *et al*, 2013) and topological properties of chromosomes (Travers and Muskhelishvili, 2005). Nevertheless, current knowledge does not allow to infer from these mechanisms a systematic decomposition of genomes into units of regulation.

Besides the conceptual problem of defining units, or modules, in a context where most entities are coupled at some level, an essential difficulty is to turn the informal notion of “functional significance” into operational methods and definitions. This notion ultimately refers to a measure of “fitness”, which is hardly accessible experimentally, given our poor knowledge of the environmental conditions under which this fitness should be evaluated. A systematic, albeit indirect approach is nevertheless possible, which uses evolutionary conservation across species as a yardstick. This approach relies on the principle that features that are shared between phylogenetically distant species reflect strong selective pressures and, therefore, functional significance. Extended to an analysis of co-evolution, this principle can reveal functionally important relationships, as now illustrated in a number of diverse problems (Junier, 2014). But although this approach could, in principle, be applied to an analysis of gene co-expression, its application is precluded by the very limited number of species for which extensive gene expression data is available.

Here, following several previous studies (Lathe *et al*, 2000; Tamames, 2001; Rogozin *et al*, 2002; Snel *et al*, 2002; Rocha, 2005; Wright *et al*, 2007; Fang *et al*, 2008), we show that synteny, the conservation of genomic contexts between phylogenetically distant bacteria, is a powerful tool which not only reveals new regulatory units but also suggests the mechanisms behind their cohesion. By systematically analyzing the conservation of proximity between orthologous genes in ∼ 1000 annotated bacterial genomes, we thus define “synteny segments” as groups of consecutive genes that are co-localized both in a particular bacterial genome and in a significant number of other, phylogenetically distant genomes. We show in the context of *E. coli* that these segments correspond to supra-operonic units with a number of remarkable structural and regulatory properties. These findings lead us to hypothesize that the most evolutionarily primitive and functionally primary modes of gene co-expression may not require any molecular factor besides a DNA and RNA polymerases: “facilitated co-transcription”, the transcription of a gene induced by the transcription of the gene located immediately upstream, sharing or not the same orientation, may be at the evolutionary origin of gene regulation and may still contribute prominently to it in current organisms. This hypothesis is supported by available RNA-seq data, leads to experimentally testable predictions, and has implications beyond bacterial genomes.

## Results

### Multi-scale genomic organization of bacterial gene expression

Microarray data provide a global view of gene expression in a given bacterial strain (Selinger *et al*, 2000; Hughes *et al*, 2000; Kapranov *et al*, 2007). This data is available for *E. coli* in the form of a compendium of 466 microarray expression profiles collected under a variety of conditions but obtained from a common platform and normalized uniformly (Faith *et al*, 2007). From this data-set, represented in Figure 1, a degree *C*_*ij*_ of co-expression between any two genes *i* and *j* can be defined, which quantifies the similarity of their profiles of activity (Alter *et al*, 2000): *C*_*ij*_ is zero in absence of correlation, positive and at most one when the two genes are expressed in similar conditions, and negative when they are expressed in different conditions (Materials and methods). Co-expression values are shown for all pairs of genes in Figure 1B, revealing a multi-scale organization of gene co-expression (Jeong *et al*, 2004; Carpentier *et al*, 2005): at the shortest genomic scales, of the order of 10 kilo-bases (kb, the scale of a single gene), small clusters of positively correlated genes are apparent (Figure 1C), while at the largest scales, of the order of 1 Mb (1/4 of the genome length), a global pattern of anti-correlation identifies two large clusters (Figure 1D). These features are conveniently recapitulated in a co-expression function Γ(*d*) (Jeong *et al*, 2004; Carpentier *et al*, 2005), defined here as the mean co-expression *C*_*ij*_ between pairs of genes separated by a given genomic distance *d*: as shown in Figure 1E (black points), Γ(*d*) presents a first decrease up to *d* ∼ 10 kb, which reflects the presence of the small correlated clusters, followed by a long plateau and a second decrease around *d* ∼ 1 Mb, which reflects the globally anti-correlated clusters.

**Figure 1:**
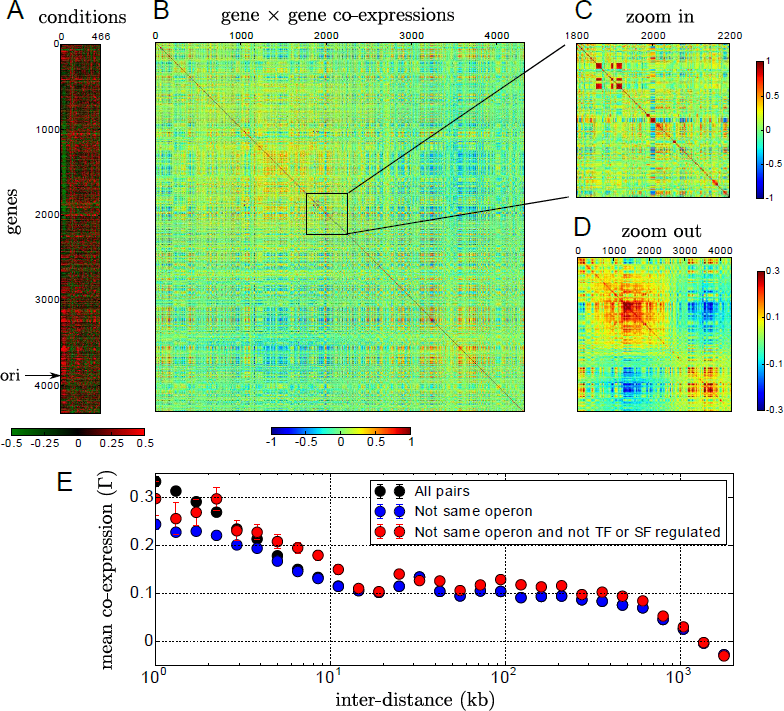
Spatial patterns of gene co-expression in E. coli. **A.** Micro-array data reporting the expression levels of 4320 genes (rows) in 466 conditions (columns) with high expression in red and low expression in green (the data is normalized so that the mean expression of a gene across conditions is zero). **B.** Matrix *C*_*ij*_ of co-expression between every pair *ij* of genes. **C.** Zoom in the co-expression of 400 genes, showing small correlated clusters of genes of size ∼ 10 kb (∼ 10 genes). **D.** Zoom out of the co-expression obtained by Gaussian filtering with a standard deviation of 10 genes, showing two globally anti-correlated clusters of size ∼ 1 Mb (∼ 1000 genes). **E.** A mean co-expression function Γ(*d*) is defined as the mean value of *C*_*ij*_ over pairs of genes at given distance *d*_*ij*_ = d. Γ(*d*) is shown when considering all pairs of genes (black dots, which are behind the blue dots when not visible), only pairs in distinct operons (blue), or only pairs in distinct operons with no known regulation by a transcription factor (TF) or a sigma factor (SF) (red).

The multi-scale organization of gene co-expression is to be related to two other hierarchical organizations: the hierarchy of regulatory mechanisms, and the hierarchy of chromosomal structures. At the bottom of these hierarchies, just above the level of genes, are the operons, which are disjoint sets of consecutive genes transcribed into a single mRNA (hereafter, we also call “operon” a gene that is transcribed in isolation); operons may also contain alternative transcription start sites (Cho *et al*, 2009) or transcriptional attenuators (Henkin and Yanofsky, 2002), implying that operonic genes can be differentially regulated (Guell *et al*, 2009). Beyond operons, several types of DNA-binding proteins coordinate gene expression. Sequence specific regulatory factors, including sigma factors (SFs) and transcription factors (TFs), enhance or repress specifically the transcription of operons (Browning and Busby, 2004). Nucleoid associated proteins (NAPs) (Thanbichler *et al*, 2005), similar to TFs but binding to a large number of non-promoter sites on the DNA, also jointly contribute to the regulation of gene expression and the structuration of chromosomes (Dillon and Dorman, 2010). At a larger scale, genes within specific genomic domains going from 10 kb to several hundreds kb are found to remain close in space (Niki *et al*, 2000; Valens *et al*, 2004; Le *et al*, 2013), suggesting the existence of micro-domains of superhelicity (Postow *et al*, 2004) that either fold into larger structures (Valens *et al*, 2004) or are themselves extended (Le *et al*, 2013). In all cases, the dynamics of theses structures depends on the cell cycle and on the growth rate of the bacterium (Kleckner *et al*, 2014); this is the case, in particular, for the two globally anti-correlated clusters apparent in Figure 1D (Supp. Figure S1; Supp. Materials and methods) (Sobetzko *et al*, 2012; Shoval *et al*, 2012).

To what extent can we explain the patterns of gene co-expressions seen in Figure 1 by these structures and mechanisms? First, the presence of operons is not sufficient to account for the short-scale patterns of co-expression whose characteristic length scale is ∼ 10 kb: considering only pairs of genes in different operons in the computation of Γ(*d*) does reduce the degree of co-expression at very short scale, but does not suppress their excess up to 10 kb (Figure 1G, blue dots). The characteristic length scale of 10 kb is, conversely, consistent with the association of these pairs of genes with supercoiled micro-domains (Postow *et al*, 2004). But even though supercoiling is known to directly affect gene expression (Peter *et al*, 2004), supercoiled micro-domains are too poorly characterized to be precisely associated with the patterns of gene expression seen in Figure 1C.

On the other hand, we observe that the main SF in *E. coli*, *σ*^70^, is associated with the global pattern of anti-correlation (Supp. Figure S1E), consistently with the known fact that most housekeeping genes expressed during exponential growth are transcribed by RNA polymerase with *σ*^70^ as subunit (Gruber and Gross, 2003). Yet, retaining only the operons known to be transcribed with *σ*^70^, and with *σ*^70^ only, does not suppress the anti-correlations (Supp. Figure S2A). A similar conclusion is reached when considering Fis, a NAP whose activity is also associated with different phases of the cell growth (Travers and Muskhelishvili, 2005) (Supp. Figure S1E). More strikingly, ignoring all genes known to be regulated by a SF or a TF leaves intact the two patterns of short and long-scale correlations (Figure 1E, red dots). In fact, the majority of correlated pairs of genes do not share a common TF or common SF (Supp. Figure S2B). The regulation of operons by SFs or TFs do not, therefore, explain the basic patterns observed in Figure 1.

Several factors beyond DNA binding proteins and local chromosomal conformations are known to affect transcription, including among several others small RNAs (Waters and Storz, 2009) and global physiological factors, which may for instance affect the concentration of available RNA-polymerases (Klumpp *et al*, 2009; Berthoumieux *et al*, 2013). None of these factors is, however, known to be associated with a characteristic length scale, and there is currently no evidence of their involvement in the chromosomal structuring of gene expression.

### Evolutionarily conserved units of synteny

Irrespectively of the nature of the underlying mechanisms, the presence of gene expression patterns that are not associated with currently known decomposition of bacterial genomes raises a simple question: How to define regulatory units beyond operons? An indirect approach to this problem is to perform an in-depth analysis of the expression data represented in Figure 1A (Ma *et al*, 2013). However, high correlations may not always reflect high contribution to the fitness of the bacterium and, conversely, moderate or low correlation may underly functionally significant relationships. Another approach consists in analyzing more directly the functional relationships by performing single or double gene knock-outs (Nichols *et al*, 2011). These perturbations are, however, too severe to identify novel regulatory units. Milder perturbations have been applied which, for instance, probe the incidence of the relative positions of genes (Block *et al*, 2012; Kuhlman and Cox, 2012; Bryant *et al*, 2015), but these experiments have not yet been performed at a scale that would permit a genome-wide analysis of regulatory properties.

### Definition of synteny segments

A generic approach to circumvent these difficulties is to rely on the principle of evolutionary conservation, i.e., to perform a comparison across multiple species with the premise that features conserved in phylogenetically distant species are under selective pressure. In absence of transcriptional data comparable to that of *E. coli* for most other bacterial strains, we apply here this approach to a systematic analysis of the conservation of relative distances between orthologous genes in different species, known as synteny properties (Lathe *et al*, 2000; Tamames, 2001; Rogozin *et al*, 2002; Snel *et al*, 2002; Rocha, 2005; Wright *et al*, 2007; Fang *et al*, 2008).

The recourse to synteny is motivated by the observation that short relative distances are under selection for co-expression (Rocha, 2005), and that gene pairs under selection extend beyond operons. Pairs of proximal genes (*d* < 10 kb) whose expression is strongly correlated in *E. coli*, including pairs of genes in distinct operons, are indeed more likely to be proximal in other phylogenetically distant bacterial strains (Supp. Figure S3). In other words, synteny can serve as a reporter of co-transcription and, hence, as a tool to infer local regulatory units beyond the operon scale. We thus analyzed, at a genome-wide level, the conservation of gene proximity by comparing the organizations of ∼ 1000 complete bacterial genomes (Materials and methods).

The first step in this analysis consists in identifying pairs of orthologous genes proximal in a significant number of bacterial genomes. Graphically, these pairs form a network, where the nodes are genes and the links represent conserved proximity between two genes (Supp. Figure S4). To analyze this network in the context of a given genome, here *E. coli*, we focus on the links that represent proximal genes in the particular genome. The resulting sub-network thus reports the pairs of genes that are both proximal in the particular genome and in a significant number of other bacterial genomes. The second step consists in defining the largest groups of genes that are both proximal in *E. coli* and in a significant number of other bacterial genomes, which correspond to the maximal sets of fully inter-connected nodes in the sub-network (Figure 2A). The genes in these groups are not necessarily consecutive in the genome of interest. We therefore finally define “synteny segments” as the segments of consecutive genes that are contained in these groups. Note that this definition does not use any information relative to the order or the orientation of the genes along the chromosome.

**Figure 2:**
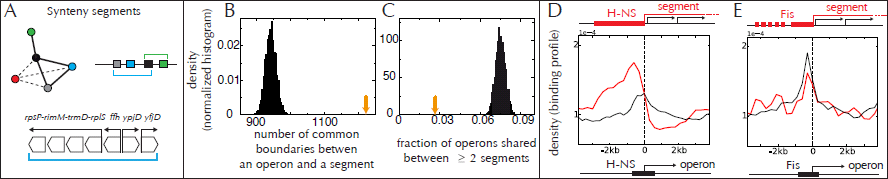
Synteny segments and their relations to structural properties of the E. coli genome. **A.** Top: Synteny relationships are recapitulated in a graph, with a link between two orthologous genes if they are proximal in a significant number of bacterial strains. The link is represented as a plain line if the genes are proximal in the particular strain of interest (here *E. coli*), and as a dotted line otherwise. The synteny segments of a particular strain are defined as consecutive genes that are all proximal both in the strain and in many others: they thus correspond to groups of genes fully connected by plain lines, as shown here for two groups that have in common the gene in black. Bottom: a synteny segment in *E. coli* containing a four-gene operon together with three single-gene operons. **B.** Distribution of the number of common boundaries between operons and synteny segments when the synteny map is translated along the genome by all possible finite numbers of genes, with the orange arrow indicating the observed value (no translation, Materials and methods). **C.** Similarly for the fraction of operons containing genes in at least two different synteny segments. **D.** The binding regions of H-NS and Fis display a statistically significant tendency of being located outside of the borders of segments (see text for p-values). **E.** Similarly, Fis shows a tendency to bind immediately outside the segments (red plain line). Contrary to H-NS, however, the resulting binding profile is not significantly different from that at the promoter of operons (black). Data from (Kahramanoglou *et al*, 2011) (Materials and methods).

Our definition of synteny segments is grounded in simple statistical principles applied to the comparison of chromosomal organizations between diverse bacterial strains. The biological relevance of these segments can be judged independently of these principles, by comparing them to structural and regulatory properties not involved in their definition.

### Synteny segments and the hierarchical structuring of chromosomes

First, we note that the 782 synteny segments found in *E. coli* (File1.txt) are distributed nearly uniformly along its genome (Supp. Figure S5), with a size distribution that follows that of its multi-gene operons (Supp. Figure S6). The same distribution is observed in vastly different genomes, suggesting that a common mechanism may be responsible for their formation (Junier and Rivoire, 2013). Next, the segments fit remarkably well within the hierarchy of known chromosome architectures. At the lowest level, their boundaries coincide in most cases with the boundaries of operons (Figure 2B). Reciprocally, operons are rarely found in two different synteny segments (Figure 2C). At a higher level, the NAPs Fis and H-NS bind preferentially at the borders of the segments (Figure 2D-E): 359 out of 444 H-NS binding regions, and 866 out of 1246 Fis binding regions, are found within 3 kb of a border (p-values 7.10^−5^ and 5.10^−6^, respectively, by comparing with translations of the operon map, see Materials and methods). In particular, we observe a clear enrichment of H-NS immediately outside synteny segments, and a depletion inside them (red profile in Figure 2D), with a staircase-like binding profile markably different from the binding profile around the promoters of operons not at a border (black profile). The same profile is obtained for the transcriptionally silenced extended protein occupancy domains (tsEPODs, of extension > 2 kb) identified in (Vora *et al*, 2009), in agreement with the fact that most of these domains overlap with H-NS binding regions and with the proposition that tsEPODs isolate supercoiled domains from each other (Vora *et al*, 2009) (Supp. Figure S7). Fis also displays a tendency for binding immediately outside of the segments with a binding profile which, however, does not significantly differ from that of operons (Figure 2E). In contrast, the highly expressed extended protein occupancy domains (heEPODs, of extension > 2 kb) also identified in (Vora *et al*, 2009) are not enriched at the border of segments; instead, they tend to be located within the segments: 102 out of the 121 heEPODs overlap with the segments (p-value 4.10^−9^).

These results show that synteny segments correspond to a level of genomic organization lying between the operons that they contain and the H-NS/Fis delimited microdomains that contain them. They are consistent with the previously proposed concepts of uber-operons (Lathe *et al*, 2000), superoperons (Rogozin *et al*, 2002), persistent genes (Fang *et al*, 2008), clusters of pathway-related operons (Yin *et al*, 2010) and cluster of statistically correlated genes (Junier *et al*, 2012).

### Internal organization of synteny segments

The relative orientation of operons contained in the synteny segments is another of their remarkable structural properties. Specifically, within synteny segments of *E. coli* made of two operons, divergent orientations are significantly over-represented while convergent orientations are under-represented (Figure 3A). Similarly, in synteny segments made of three operons, patterns of divergence and co-directionality are over-represented, whereas patterns of convergence are under-represented. These features are shared across other bacterial strains. A decomposition into operons is not available in most cases, but we can circumvent this limitation by comparing the relative number of co-directional, divergent and convergent successive genes, inside versus outside synteny segments (Figure 3B). We thus observe that the ratio of divergent over convergent orientations of genes, which must be 1 over an entire circular chromosome, is larger inside than outside synteny segments for virtually all bacterial genomes; genomes that do not share this property consist, without exception, of > 90% co-directional gene pairs. Similarly, the ratio of co-directional over divergent or convergent orientations is systematically larger for successive genes inside synteny segments (Figure 3B).

**Figure 3:**
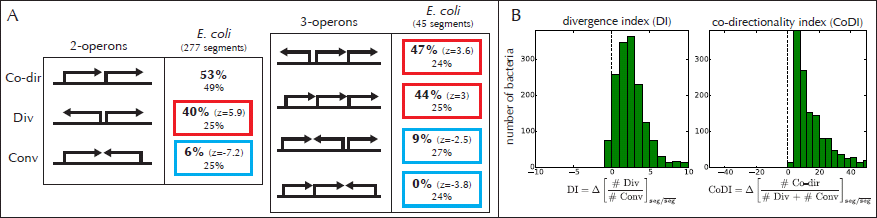
Gene orientations within synteny segments. **A.** Statistics in *E. coli* over all synteny segments made of 2 and 3 operons exactly, showing that some organizations are over-represented (red squares, with *z* > 1.65 (*P* < 0.05) indicating standard deviation against a randomized set) or under-represented (blue squares, *z* < −1.65). Percentages on the first and second line respectively correspond to statistics within the segments and overall, for the total of 2647 operons. **B.** For each genome, we compute a divergence index (DI) defined as the difference between the ratio of divergent over convergent orientations, which must be 1 over the entire chromosome, inside versus outside segments. We also compute a co-directionality index (CoDI) defined as the difference of the ratio of co-directional over divergent or convergent orientations. The green histograms show the statistics of these indexes over our dataset of ∼ 1000 annotated bacterial strains.

### Synteny segments and co-transcription

Beside these remarkable structural properties, gene expression data provides direct support in favor of the hypothesis that synteny segments are the evolutionary imprint of fundamental genomic units of co-regulated genes, which may consist of several operons and whose co-regulation may not require that they all share the same SF or TF.

### Co-transcription within and between synteny segments

Intra-segment co-expression indeed occurs at high levels, as shown in Figure 4A using the same co-expression function Γ(*d*) as in Figure 1E (the black dots are identical in the two figures). Specifically, pairs of genes in a same segment (in green) display, at short distances (*d* < 5 kb), a level of co-expression that is comparable to pairs in a same operon (in red) and, at higher distances (5 kb < *d* < 10 kb), an even higher level. Excluding intra-operon pairs (and segments < 10 kb, which contribute only at short distances) clearly indicates that intra-segment correlations are not just due to intra-operon correlations (blue dots in Figure 4B). TFs and SFs are also seemingly irrelevant: the pairs of genes from different operons with no TF and with different SFs display the same level of co-expression (yellow dots).

Together with the particular orientations of operons within segments, these observations suggest that synteny segments represent supra-operonic transcriptional units controlled by a subset of “entry points”, with co-transcription of divergent operons and of successive co-directional operons. As a result, a same TF may regulate several operons within a same unit. This proposition implies that larger segments may require less TF per operon than smaller segments, which is indeed supported by the data (Figure 4C). It is also consistent with the known associations between the patterns of orientation and the regulation of operons by TFs (Korbel *et al*, 2004; Warren and ten Wolde, 2004; Hershberg *et al*, 2005).

**Figure 4:**
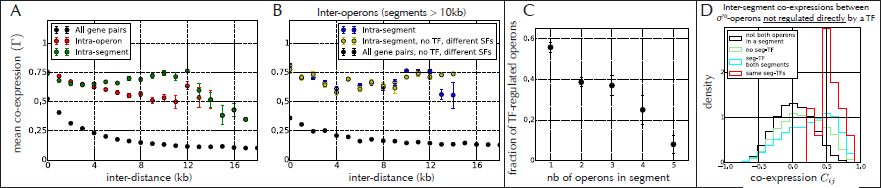
Intra-segment co-expression. **A.** As in Figure 1E, the black dots represent Γ(*d*), the mean value of *C*_*ij*_ for pairs of genes at a given distance *d* along the chromosome of *E. coli.* The red and green dots represent the same quantity but restricted, respectively, to pairs of genes in the same operons and in the same synteny segment. **B.** Similar to A, but now excluding pairs of genes in a same operon and retaining only segments larger than 10 kb. In blue, pairs of genes in a same synteny segment but in distinct operons. In yellow, adding the constraints that the two genes are in operons that are not regulated by any transcription factor (TF) and not regulated by a common sigma factor (SF). The high level of correlation, independent of the inter-operon distance, is observed irrespectively of the relative orientation, and holds both for divergent and for co-directional operons (Supp. Figure S8). It also holds for small segments < 10 kb, even though the average co-expression level is lower (Supp. Figure S8). **C.** Fraction of operons controlled by a transcription factor (TF) as a function of the number of operons in the segment, showing that operons in large segments tend to be less TF-regulated in average. **D.** Distribution of the co-expression *C*_*ij*_ between different classes of pairs of genes, all taken to be regulated by the SF *σ*^70^ but not directly by any TF: in black, pairs with at least one gene not in a segment; in green, operons in segments but with no seg-TF (i.e., not belonging to a segment containing a TF-regulated operon); in cyan, operons with seg-TF; in red, operons with same seg-TF.

The proposition that promoter-specific regulators such as TFs and SFs do not act exclusively on operons, but more broadly on segments, leads us to introduce the notion of “seg-TF”: a TF is a seg-TF for a gene if it is a TF for an operon in the same segment (a “seg-SF” is defined similarly). A gene may thus have a seg-TF but no TF of its own. The relevance of this notion is demonstrated in Figure 4D: pairs of genes regulated by the SF *σ*^70^ but in different segments and with no TF of their owns are significantly more co-transcribed when they have exactly the same seg-TFs (red distribution, to be compared with the cyan distribution, where the genes have different seg-TFs). Interestingly, pairs of such genes are also significantly more co-transcribed when they both have a seg-TF (irrespectively of its identity, cyan histogram) than when they both have no seg-TF (green histogram), suggesting an indirect contribution from the regulation of TFs by other TFs (Ma *et al*, 2004).

Common seg-TF regulation thus implies long-range co-expression. Yet, segments with exactly the same set of seg-TFs are few, which suggests that this phenomenon plays, overall, a secondary role (Supp. Materials and methods; Supp. Figure S9). As our analysis relies exclusively on the evolutionary conservation of gene proximity, it does not directly address the problem of long-range co-regulation. Remarkably, however, it points towards an evolutionary link between short and long-range co-transcription. Indeed, pairs of genes that are distant in a genome, but in synteny in other genomes, are in average more co-expressed than those not in synteny (Supp. Figure S10). This phenomenon appears to be specific, in the sense that it does not apply to immediately neighboring genes (Supp. Figure S10). It suggests that operons that were previously proximal but later set apart evolve, or have evolved, similar cis-regulation (Wang *et al*, 2011).

### Mechanisms of co-transcription (or lack thereof)

What are the molecular mechanisms behind the co-transcription of distinct operons within a same segment? We propose here a parsimonious explanation that we call “facilitated co-transcription”: in absence of additional molecular factors or specific inter-gene sequence motifs, the transcription of an operon is facilitated by the transcription of the operon located immediately upstream. In other words, the mere interaction of RNA polymerases with DNA may constitute the basic mechanism by which operons within a same segment are co-transcribed.

Facilitated co-transcription may have different origins, depending on the relative orientation of the genes. For co-directional genes, it may be caused by “transcriptional read-through”, the transcription of consecutive operons by RNA polymerases overriding the signals of termination (Henkin and Yanofsky, 2002), which is indeed known to be a major source of transcripts in bacteria (Wade and Grainger, 2014). For divergent genes, evidence from eukaryotic genomes indicates the presence of a polymerase at a promoter is associated with high likelihood with the presence of another polymerase engaged upstream and in the opposite orientation (Core *et al*, 2008). The pervasive antisense transcription thus induced at bidirectional promoters, first reported in eukaryotes (Xu *et al*, 2009), is also found in bacteria (Dornenburg *et al*, 2010; Lasa *et al*, 2011). In this case, the underlying physical mechanism may involve supercoiling: the torsional stress induced by transcribing RNA polymerases is indeed known to enhance or repress the transcription of nearby genes (Liu and Wang, 1987; Hatfield and Benham, 2002; Meyer and Beslon, 2014). Increases in the local concentration of RNA polymerases may also play a role (Junier, 2014). The co-transcription of operons within a segment may thus not require any specific mechanism.

To support the hypothesis of facilitated co-transcription and shed light on the underlying mechanisms, we consider strand-specific RNA expression profiles obtained by RNA-seq (Core *et al*, 2008). A first observation, consistent with previous reports of wide-spread pervasive transcription (Wade and Grainger, 2014), is that transcription of a gene is associated with a proportional transcription of its anti-sense sequence, independently of whether the gene belong to a segment and, independently of the orientation of the upstream gene (Supp. Figure S11). To assess the role of transcriptional read-through, we consider pairs of consecutive and co-directional operons and examine the correlation between the transcription of the first gene in the second operon and the transcription of the upstream, non-coding inter-operonic sequence (see small drawing on top of Figure 5A). The two are indeed strongly correlated, in agreement with wide-spread transcriptional read-through; remarkably, the correlation is larger when the two operons are in a same segment (Figure 5A). Given the general correlation between sense and anti-sense transcription (Supp. Figure S11), anti-sense transcription can be expected to play a major role in the co-transcription of consecutive divergent operons. This is indeed corroborated by the observation of a systematic enrichment, when the gene is in a segment, of the correlations between the transcription of a gene and the anti-sense transcription of the upstream non-coding inter-operonic sequence (Figure 5B).

**Figure 5:**
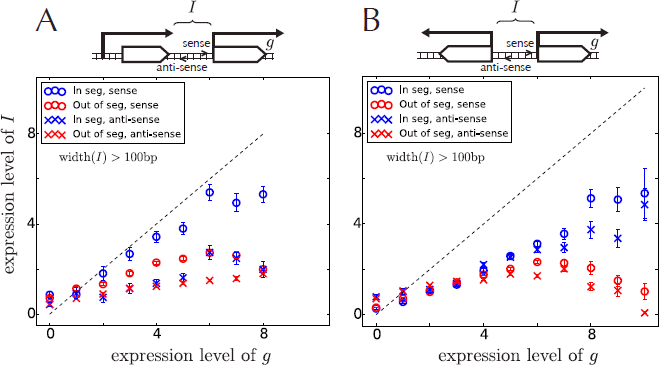
Evidence for facilitated co-transcription. **A.** For pairs of consecutive co-directional operons, correlation between the transcription of the first gene *g* in the second operon and the sense or antisense transcription of the upstream inter-operonic sequence *I* (small drawing on top). Sense transcription of *I* (circles) correlates strongly with the transcription of *g*, much more than anti-sense transcription (crosses), and this correlation is stronger for operons inside a same segment (in blue). These observations are consistent with widespread transcriptional read-through, particularly inside segments. Here, we consider inter-operonic regions longer than 100 bp, hence excluding the possible presence of mis-annotated operons (McClure *et al*, 2013) **B.** For pairs of consecutive divergent operons, correlation between the transcription of one of the two genes (*g*) and the transcription of the upstream sense or anti-sense inter-operonic sequence (*I*). The transcription level of *I* is lower than the anti-sense transcription of *g* (Supp. Figure S11) and, compared to A, there is no significant difference between the sense and anti-sense transcription of *I*. We observe, however, a higher level of transcription of the inter-operonic region *I* when genes belong to segments. These plots were made using the RNA-seq data from (McClure *et al*, 2013) with transcription start sites retrieved from RegulonDB, by considering for each gene the furthest upstream start sites identified in the Morett dataset (Salgado *et al*, 2012) (Materials and methods).

From these observations, we conclude that facilitated co-transcription, in the form of transcriptional read-through and anti-sense co-transcription, is a wide-spread phenomenon, particularly occurring inside segments. This provides a rationale for the high level of co-transcription within segments, including in the absence of TF (Figure 4). Further insights could be gained from new experiments, using for instance synthetic constructs comprising two operons with varying inter-operonic sequences and inserted at different locations in a bacterial genome.

## Discussion

By systematically comparing the relative positions of orthologous genes in multiple bacterial genomes, we identified “synteny segments”, corresponding to groups of consecutive genes in one genome that are also proximal in a significant number of other genomes. Consistently with previous studies (Lathe *et al*, 2000; Rogozin *et al*, 2002; Fang *et al*, 2008; Yin *et al*, 2010; Junier *et al*, 2012), these synteny segments fall within the known hierarchy of genomic structures: they contain and extend the operons, and are contained within the micro-domains defined by H-NS and Fis binding sites. Inside a segment, operons are most often co-directional or divergent, and genes are co-expressed at a comparable Level, whether in a same operon or not. In particular, two genes can be co-expressed in the absence of a direct regulation by a SF or a TF, just by being proximal and aligned either co-directionally or divergently. This very mechanism also allows for a long-range co-regulation of genes belonging to distinct segments, when the segments share common TFs – what we termed “seg-regulation”. Taken together, these findings identify the synteny segments as evolutionary signatures of supra-operonic units of co-transcribed genes.

### Facilitated co-transcription as a primitive mode of regulation?

We proposed that facilitated co-transcription, the facilitated transcription of a gene by the transcription of the gene immediately upstream, in absence of additional molecular factor or specific inter-genic sequence motif, constitutes a primary form of regulation in current bacteria (Figure 6A). We extend here this hypothesis by proposing that facilitated co-transcription also represents its most primitive form, when considering the problem from an evolutionary standpoint.

**Figure 6:**
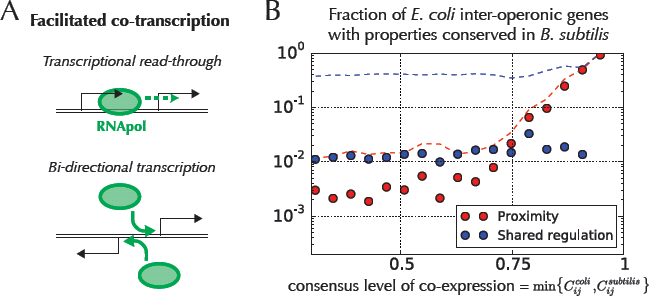
Facilitated co-transcription and evolution. **A.** Facilitated co-transcription, the facilitated transcription of a gene by the transcription of the gene immediately upstream, may take two forms: transcriptional read-through, when two neighboring genes are co-directional, and anti-sense co-transcription, when their orientations are divergent. **B.** Fraction of pairs of genes *ij* in different operons in *E. coli* that are proximal (< 20 kb, red dots) or regulated by a common TF or SF (blue dots) both in *E. coli* and in *B. subtilis*, as a function of the minimal value of the co-expression *C*_*ij*_ between the two strains, 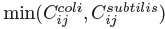. A high value of 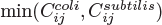 indicates a conservation of co-expression. For comparison, the dotted lines represent the same quantities when considering properties, proximity or shared regulation, in *E. coli* only. The conservation of co-expression thus appears to be associated with a conservation of proximity rather than with a conservation of regulatory factors.

In this scenario, gene clustering would have came first and TF-specific regulation would represent a relatively recent and secondary addition, tailored to the need of each specific lineage. In support to this view, we note, first, that TFs and their network have indeed been shown to evolve quickly compared to other genetic networks (Babu *et al*, 2006; Shou *et al*, 2011), and, second, that the clustering of genes encoding different major fundamental processes, such as replication and translation, is highly conserved throughout evolution (Junier, 2014). Besides, the rewiring of gene regulatory networks shows only a marginal impact on both the genome-wide transcription and the fitness of *E. coli* (Isalan *et al*, 2008). Next, and consistently with an evolution of TF-regulation on top of a pre-existing regulation by transcriptional read-through, we observe that among operons of *E. coli* preceded by a co-directional operon, only 307 are regulated by a TF while 992 are not, a difference that is highly significant (p-value 10^−7^). We also note that only 10% of the operons are annotated with terminators (25% when including more complex transcription attenuation) (Salgado *et al*, 2012). Another line of evidence comes from analyzing the conserved features of genes that are strongly co-expressed in phylogenetically distant bacteria. A comparison between *E. coli* and *B. subtilis*, two strains for which extensive transcriptional data is available (Faith *et al*, 2007; Nicolas *et al*, 2012), reveals that the pairs of genes with conserved co-expression typically correspond to genes that are proximal in the two genomes, rather than to genes that are controlled by a common TF (including non-orthologous ones; Figure 6B). Along the same line, while transcription is known to be regulated at a global level by supercoiling (Blot *et al*, 2006), with a demonstrated influence on fitness (Crozat *et al*, 2005, 2010), deleting Fis, one of the NAPs which with H-NS controls supercoiling, has only marginal effects, depending on the conditions under which the bacterium grows (Crozat *et al*, 2010) As TFs acting at promoter sites, but at a more global level, NAPs may thus only modulate the more fundamental patterns of co-expression imposed by the relationships of proximities between genes (Cameron *et al*, 2011).

The hypothesis that facilitated co-transcription of co-directional and divergent genes is at the evolutionary origin of gene clustering also disposes of the paradoxes usually associated with the evolution of operons. The selfish operon scenario has indeed challenged the commonly-hold assumption that selection for co-regulation drove the evolution of operons (Lawrence and Roth, 2002). In particular, it has questioned the selective advantage of evolutionary intermediates when forming a new operon by bringing together several genes and an operator. Under our hypothesis, the clustering of transcriptionally independent genes may enhance their co-expression, independently of the presence of operators. This may confer an adaptive benefit to the bacteria before an operon is formed. In agreement with this scenario, gene clustering is found to be under positive selection (Junier and Rivoire, 2013). The selfish operon scenario, which proposes horizontal gene transfer as driving the formation of operons, is not, on the other hand, supported by evidence (Pál and Hurst, 2004; Price *et al*, 2005).

Finally, let us point out that the conservation of gene proximity extends beyond bacteria, to the eukaryotic organism *S. cerevisiae*: orthologs of genes in synteny in bacteria that are proximal in *S. cerevisiae* display, indeed, a high level of co-transcription (Figure S12). From the point of view of regulation, microarray data associates virtually every gene of *S. cerevisiaeto* at least one TF but ChIP-seq data suggests that only a small fraction of these associations stem from direct physical interactions (Geistlinger *et al*, 2013). Higher than expected levels of co-expression between proximal genes have then been attributed to chromatin remodeling (Batada *et al*, 2007). Facilitated co-transcription offers an alternative explanation, without, however, excluding additional factors. More generally, clusters of co-expressed genes are a common feature of eukaryotic genomes (Michalak, 2008). As these genomes do not contain operons and have regulatory mechanisms significantly different from those of bacteria, the presence and conservation of gene clustering further support the hypothesis of a generic mechanism causing the co-transcription of proximal genes. Transcription read-through and divergent promoters have, in fact, also been proposed to account for the conservation of gene cluster in mammals (Semon and Duret, 2006) and supercoiling is recognized as a crucial factor for the local properties of gene regulation (Kouzine and Levens, 2007).

The hypothesis that facilitated co-transcription is both primitive and primary still leaves open two basic evolutionary questions. First, it shifts the challenge from explaining how gene expression became coupled to the challenge of explaining how it became uncoupled (Singh *et al*, 2014). While this problem is beyond the scope of the present work, we note that transcriptional termination is as regulated as initiation (Henkin and Yanofsky, 2002), and can be strongly conserved (Merino and Yanofsky, 2005), although the extent to which inter-operon transcriptional termination is regulated remains to be characterized. We also note that eukaryotic chromosomes have evolved elaborated mechanisms to keep genes in a repressed state. Second, the selective force behind the formation of operons remains unclear, if, as suggested by the data, proximal genes can be co-expressed at comparable level whether they are in a same operon or not. Among non-exclusive possibilities, one may invoke the control of the production of unnecessary proteins (Kovács *et al*, 2009) or the maintenance of a precise stoichiometric ratio between the different operonic proteins (Junier, 2014).

### Towards a systematic identification of regulatory units

Given the difficulty to clarify the nature and relative contribution of different regulatory mechanisms, identifying regulatory units by making minimal assumptions on the possible mechanisms is critical (Michalak, 2008). Our analysis of synteny rests essentially on a single assumption: conservation across multiple species reflects strong selective pressures, and, therefore, important functional constraints (Mering, 2003). This principle is general and applies beyond synteny, to the analysis of the co-occurrence of genes or to the evolution of their amino acid sequences. In these different contexts, it has also shown its value by revealing functional features that studies focusing on a single system or species had overlooked (Junier, 2014).

Approaches based on evolutionary conservation, being statistical in nature have, however, their own limitations. The synteny segments that we identify are, therefore, not expected to always coincide with “true” regulatory units. Several factors are preventing an exact correspondence: (i) some properties of co-regulation may be strain-specific and therefore not evolutionarily conserved; (ii) the comparison of genome architecture between strains relies on the identification of orthologous genes, which is imperfect and leaves out many orphan genes (Yamada *et al*, 2012); (iii) the number of synteny relationships that we can detect is a function of the number of available genomes, and given the total number of gene pairs (∼ 10^7^), disposing of only ∼ 10^3^ genomes sets a strong statistical constraint; (iv) the genomes are phylogenetically related and a conservation of relative distance between genes of closely related strains may reflect insufficient time to divergence, rather than common selective pressure. The first three factors can cause us to overlook many co-regulated genes. The last factor, phylogenetic bias, implies instead that we may overestimate the number of co-regulated genes. To mitigate this phylogenetic bias, we relied on a simple approach that proved its value in other contexts (Morcos *et al*, 2011). The properties of the synteny segments relative to the location and orientation of operons, to the binding sites of NAPs and to transcriptional data indicate that, despite these limitations, the approach reveals non-trivial features. In light of our results, we therefore expect that improved identification of orthology, exploitation of a larger number of genomes and more sophisticated corrections of the phylogenetic bias should have the potential to refine the identify of supra-operonic regulatory units as synteny segments, and thus to reveal further information on their nature and evolution.

## Materials and methods

### Co-expression analysis in *E. coli*

We use the microarray expression profiles from the M^3D^ database (Faith *et al*, 2007) to define the activity *x*_*si*_ of gene *i* in condition *s*, by averaging the values associated with the probes overlapping with gene *i*, and subtracting the mean expression across conditions, so that ∑_*s*_ *x*_*si*_ = 0 for all *i*. From these profiles, we define the co-expression of a pair *ij* of genes as the correlation matrix 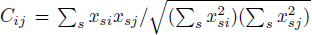. The autocorrelation Γ(*d*) is defined as the average value of *C*_*ij*_ over the pairs *ij* of genes at a given distance *d*±Δ*d*, i.e., 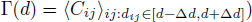, with Δ*d* = 0.5 kb.

### Genomes and orthology

Sequenced bacterial genomes were downloaded from NCBI, yielding an initial data set of *M*_0_ = 1432 genomes annotated in terms of *N* = 4467 clusters of orthologous genes (COGs). COGs are defined on the principle that any group of at least three genes from distant genomes that are more similar to each other than to any other genes from the same genomes should belong to the same COG (Tatusov *et al*, 2000); as a result, a genome may contain one, several or no gene associated with any given COG. We further removed genomes with size below 500 kb or with less than 60 % of genes annotated by COGs to obtain the *M* = 1108 genomes used in our analysis.

### Identification of synteny segments

We first identify all pairs of orthologous genes that tend to remain proximal in a significant number of genomes, and gather them in a graph (network of synteny relationships, Supp. Figure S4). To this end, for each pair of COGs *ij*, we define its relative distance in a genome as the minimal distance, in base pairs, between its respective genes – distance between two genes is measured in base pairs, from the mid-point of their nucleotide sequences. The distribution of this distance across all genomes is then computed by taking into account the phylogenetic biases coming from the uneven sampling of the space of genomes, which leads to the definition of an effective number of genomes *M*′ < *M*, with here *M*′ ≃ 470 (Morcos *et al*, 2011) (Supp. Materials and methods). We then assign a *p*-value 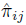) to the pair *ij* by comparing this distribution with the distribution obtained from a null model where genes are distributed independently and uniformly across *M*′ genomes (Supp. Figure S13).

**Figure S1:**
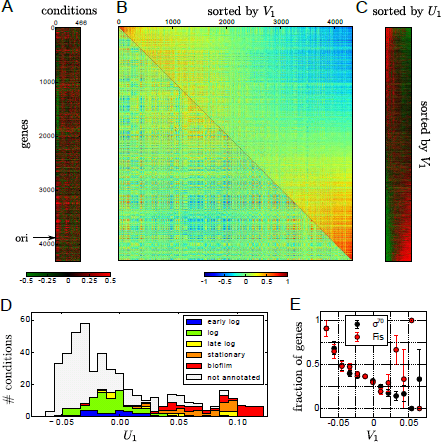
**A.** As in Figure 1A, micro-array data reporting the expression levels of 4320 genes (rows) in 466 conditions (columns) with high expression in red and low expression in green. **B.** Applying a singular value decomposition to the micro-array data yields two principal components, *V*_1_ along the genes and *U*_1_ along the conditions. The co-expression matrix of Figure 1B is shown here with, above the diagonal, the genes sorted by *V*_1_: this component classifies the genes according to their contribution to one of the two anti-correlated clusters visible in Figure 1D. **C.** Same expression data as in A, but with the conditions sorted by *U*_1_ and the genes sorted by *V*_1_, thus revealing the main pattern of variation. **D.** Distribution of the conditions along the principal component *U*_1_, with different colors for the different phases of growth at which the measurements of transcriptional activity were made, showing that *U*_1_ correlates with the growth rate. **E.** Fraction of genes controlled by *σ*^70^ (black) and with a binding site for the NAP Fis (red) as a function of *V*_1_, showing that genes that are transcribed in growing phases (negative values of *V*_1_) are more likely to be regulated by *σ*^70^ and bound by Fis.

**Figure S2:**
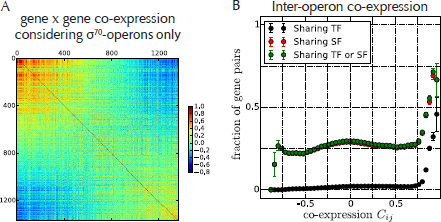
**A.** Co-transcription between the 1231 genes having *σ*^70^ as unique SF. Genes are reordered along the first component *V*_1_ from SVD decomposition of the data as in Figure S1B. **B.** Fraction of pairs of genes belonging to different operons that share a TF, a SF or one of the two, showing that, except at very high level co-expression (*C*_*ij*_ > 0.85), the majority (∼ 75%) of correlated pairs of genes do not share a common TF or SF.

**Figure S3:**
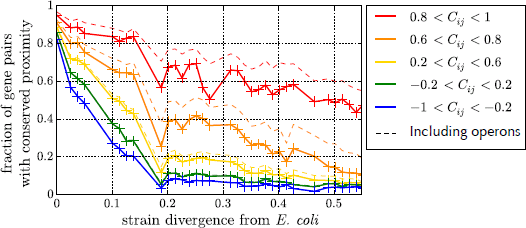
*Proximal genes in E. coli tend to be proximal in other bacterial strains if co-expressed.* Here, two genes are considered as proximal when their relative distance is below 10 kb, and strain divergence is defined as global sequence divergence (see Supp. Materials and methods). The plain lines exclude pairs of genes in a same operon in *E. coli*, while the dotted lines include them. This graph shows that the relation between the co-expression of genes and the evolutionary conservation of their proximity (Rocha, 2005) extends beyond operons.

**Figure S4.**
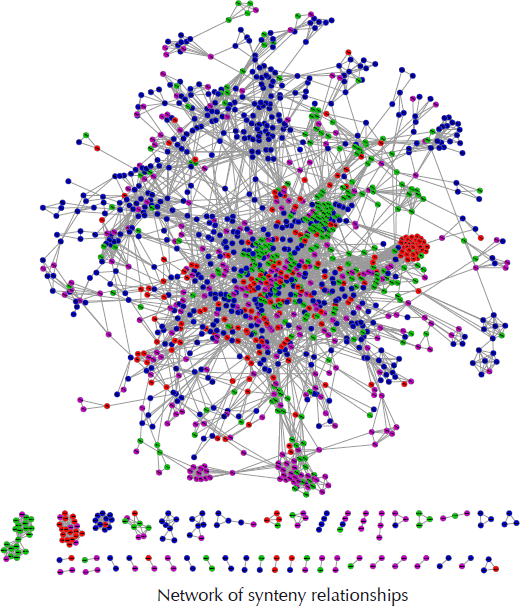
Network of synteny where a link is present between two orthologous genes if the two genes are proximal in a significant number of strains. Here, for the sake of clarity, only a subset of the full network, comprising 1455 orthologous genes (annotated into COGs), is shown. Colors indicate four functional classes of COGs: red for information storage and processing, green for cellular processes and signaling, blue for metabolism and purple for poorly characterized.

**Figure S5.**
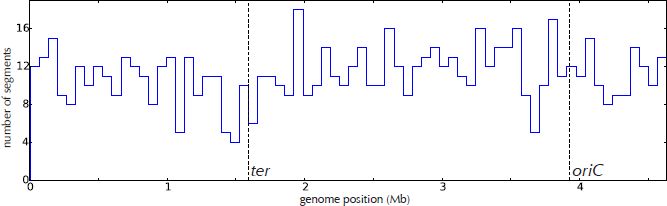
Genomic distribution of segments in *E. coli*: the histogram of the location of the segments along the chromosome reveals a fairly uniform distribution (bin size of 65 kb). The vertical dashed lines indicate the origin (*oriC*) and terminus (*ter*) of replication.

**Figure S6.**
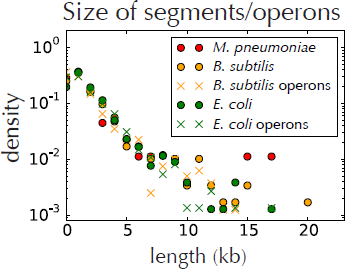
Size distributions of synteny segments (solid circles) and multi-gene operons (crosses) in three phylogenetically distant bacteria, showing a similar exponential decrease up to ∼ 10 kb. For *Mycoplasma pneumoniae*, no operon map is currently available (Guell *et al*, 2009).

**Figure S7.**
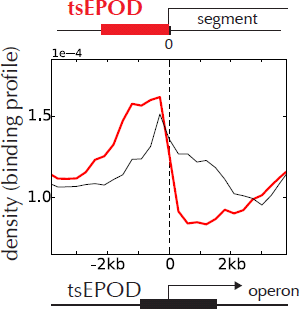
Binding profile of tsEPODs (Vora *et al*, 2009) with respect to synteny segments (red plain line) and operons (black), showing, as in the case of H-NS (Figure 2D in main text), a strikingly high density of tsEPODs at the external boundaries of segments together with a depletion inside segments. In agreement with their role in transcription silencing (Dorman, 2007), we also observe an enrichment around the promoter region, and over the first gene for operons not at the border.

**Figure S8.**
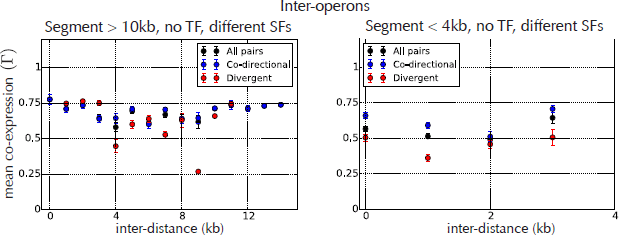
**A.** The black points correspond to the yellow points in Figure 4B, and the blue and red points show that restricting to co-directional or divergent pairs has little incidence. **B.** Similar to A, but considering the smallest segments (< 4 kb) instead of the largest ones (> 10 kb): the overall level of the correlation is lower for shorter segments.

**Figure S9.**
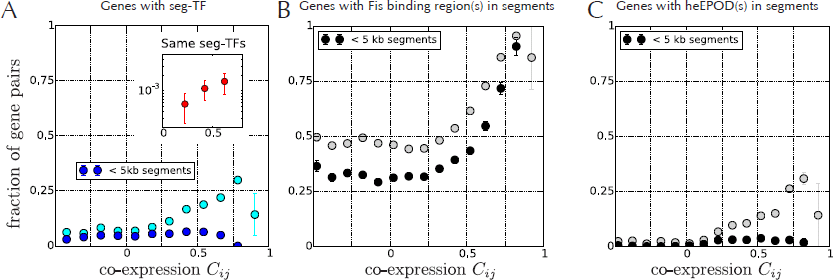
*Inter-segment co-expression.* **A.** Cyan: fraction of gene pairs with a seg-TF as a function of their correlation. Blue: considering only genes that belong to segments below 5 kb. Inset: fraction of gene pairs with same set of seg-TFs as a function of their correlation. **B.** Gray: fraction of gene pairs with a Fis binding region in the segment to which they belong. Black: considering only genes in segments below 5 kb. **C.** Fraction of gene pairs with a heEPOD (Vora *et al*., 2009) in the corresponding segment as a function of the level of co-expression *C*_*ij*_. Black: considering only genes in segments below 5 kb.

**Figure S10.**
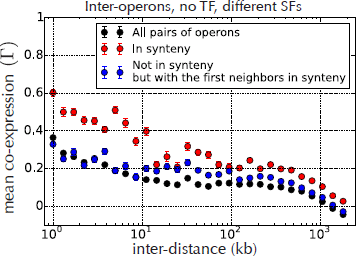
Co-expression between *E. coli* genes in different operons that are not regulated by any TF and that do not share the same SF (black points). In red, we restrict to pairs that are in synteny, independently of whether they are proximal in the chromosome of *E. coli*: these pairs are in average more co-expressed than those not in synteny. The phenomenon appears to be specific since replacing the first gene in these pairs by its nearest neighbor not in synteny (while keeping the second gene) significantly decreases the level of co-expression at all distances.

**Figure S11.**
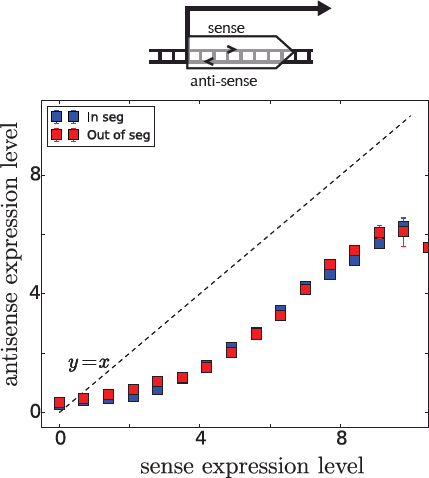
The level of anti-sense transcription is systematically correlated with the level of sense transcription, independently of whether the gene belongs to a segment or not, suggesting pervasive promiscuous transcription. (RNA-seq data as in Figure 5).

**Figure S12.**
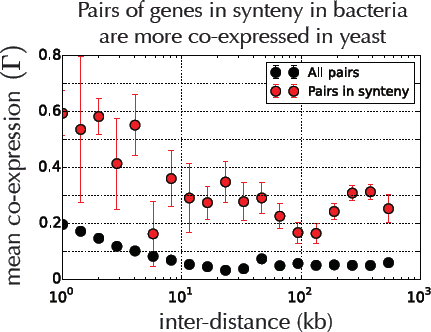
This figure is the counterpart of Figure 5A for *S. cerevisiae*, using again transcription data from the M^3D^ database (Faith *et al*., 2007) and the same definition of synteny, which is based on bacterial genomes and therefore does not include any yeast genome. The black dots represent the mean co-expression averaged over all pairs genes, and the red dots over the pairs of genes in synteny in bacterial genomes (pairs of genes in different chromosomes are not considered).

**Figure S13.**
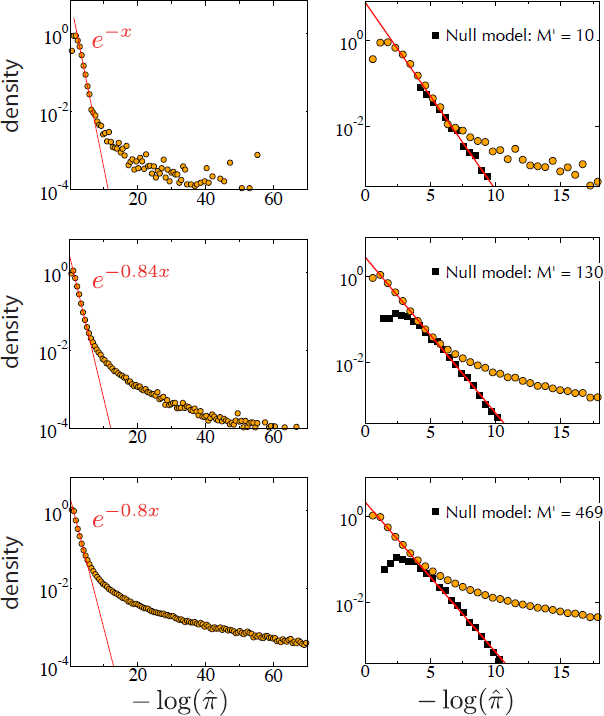
Probability density of 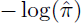 for the empirical data (orange circles) at three phylogenetic depths corresponding to three effective number of genomes: *M* = 10,130,469 (we consider the latter case for the study). Left panels: For small enough values of 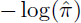, the density decays exponentially with 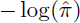 (red lines). The deviation from an exponential at large values indicates the conservation of co-localization. Right panels: For the null model where gene positions are randomized (black squares, right panels), with as number of genomes the effective number *M*′ corresponding to *δ* (*M*′ = 10,130,469, respectively), the exponential decay extends to larger values of 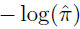.

Given the large number of pairs *ij* under study (∼ 10^7^), some of the *p*-values 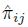 are borne out to be very small, even under the null model. One more step is therefore required to set a threshold of significance for these *p*-values. This is achieved by comparing the empirical distribution *f*(*π*) of 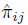 with its distribution under the null model, *f*_0_(*π*). The fraction of false positives when calling significant the pairs *ij* with 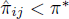 can be estimated from the ratio of the areas below these curves, as 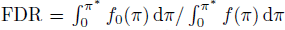. A given false discovery rate FDR (the fraction of false positives given *π*\*), here taken at 5% (more stringent values lead to similar results; see e.g. (Junier and Rivoire, 2013), thus selects a threshold of significance *π*\* (Benjamini and Hochberg, 1995).

The synteny segments of a given genome are eventually given by the maximal sets of genes that are fully interconnected in this network of synteny and that are close-by along the genome (Figure 2A); two genes are considered to be close-by if they are separated by less than 50 kb, or less than the distance that is associated to their statistical tendency of proximity (Supp. Materials and methods; taking a smaller value than 50 kb does not affect the results).

### Operons and protein binding regions

The annotations in terms of operons, TFs and SFs were all taken from RegulonDB (Salgado *et al*, 2012). H-NS and Fis DNA-binding data were retrieved from (Kahramanoglou *et al*, 2011) and we consider the binding sites obtained in the early exponential phase – similar results are found for the other phases for H-NS (binding sites are not detected for Fis in the stationary phase). Extended protein occupancy domains (EPODs) were retrieved from (Vora *et al*, 2009).

### Statistical significance of the synteny segments with respect to the hierarchical organization of chromosomes

To assess the significance of the relations between synteny segments and operons (Figure 2B-C), we consider randomizations of the data where the synteny map is translated with respect to the gene map by *m* genes, for all 4144 possible values of *m*. To assess the significance of the relations between synteny segments and binding regions of proteins (Fis, H-NS, tsEPODs, heEPODs), the synteny map is translated with respect to the operon map.

### RNA-seq data

RNA-seq data were retrieved from (McClure *et al*, 2013) under the form of .sra files. RNA reads were mapped to the genome of *E. coli* K12 MG1655 using bowtie2. The number of reads per bp was then computed as the genomic coverage of the data (using genomeCoverageBed and the flags “-d-split”), with the final expression levels equal to the log-value of the mean number of reads found in the regions of interest (*I* or *g* in Figure 5). We considered datasets for which more than 90% of the reads were uniquely mapped. Results in Figure 5) are thus the average of six different conditions corresponding to the following GEO Accession Number: GSM1104381 (sgrS- with vector), GSM1104384 (sgrS- with sgrS+ plasmid), GSM1104387 (WT in LB +*α*MG), GSM1104402 (WT in defined medium with glycerol −*α*MG), GSM1104405 (sgrS- in defined medium with glycerol +*α*MG) and GSM1104408 (sgrS- in defined medium with glycerol −*α*MG).

### Data access

The list of synteny segments is provided in File1.txt.

## Acknowledgements

We thank Olivier Espeli, Luis Serrano and Eva Yus for interesting discussions. This work was supported by a Novartis grant (to I.J.) and by ANR grant ’CoevolInterProt’ (to O.R.). The research Leading to these results has received funding from the European Research Council under the European Union’s Seventh Framework Programme (FP7/2007-2013) / ERC grant agreement n609989.

## 1 Supplementary Materials and methods

### 1.1 Relation between gene co-expression and growth conditions

The two globally anti-correlated clusters seen in Figure 1D can be interpreted functionally by relating them to the conditions under which the genes are expressed. To this end, a statistical method extending principal component analysis, known as “singular value decomposition”, can be applied, which reorders the genes and the conditions in a consistent way, according to their main axes of variation (Alter *et al*., 2000). Specifically, the singular value decomposition of the transcription profile matrix *x*_*si*_ is of the form *x*_*si*_ = ∑ *ρ*_*k*_*u*_*sk*_*v*_*ik*_, with *ρ*_1_ ≥ *ρ*_2_ ≥ … ≥ 0 the set of singular values. {*u*_*s*_}_*s*=1…#conditions_ and {*v*_*i*_}_*i*=1…#genes_ are, respectively, orthonormal basis of the gene space and of the condition space. The top singular vectors *U*_1_ and *V*_1_ have components (*U*_1_)_*s*_ = *u*_*s*1_ and (*V*_1_)_*i*_ = *v*_*i*1_, and define the main axes of variation in the gene space and condition space, respectively.

As a result, we obtain an ordered list of genes with the most anti-correlated genes at the two extremes, and an ordered list of conditions depending on whether they induce one or the other set of genes (Figure S1B-C). These lists indicate a simple interpretation of the two globally anti-correlated gene clusters in terms of phase of cell growth. Indeed, one gene cluster is preferentially expressed during exponential growth and the other during stationary phase (Figure S1D). This association of different growth rates with different overall patterns of gene expression is well recognized (Shoval *et al*., 2012). The preferential location of the anti-correlated genes on different halves of the genome is consistent with previous analyses (Sobetzko *et al*., 2012).

### 1.2 Measure of sequence divergence

For computing genome weights (*w*_*g*_, see below) and in Figure 3, the sequence divergence *δ* between any two genomes is computed as *δ* = 1 − *f*, where *f* is the average fraction of common amino acids between a selection of 10 genes. The 10 selected genes are associated with the COGs 126G, 173J, 202K, 2255L, 481M, 497L, 541U, 544O, 556L, 1158K. These COGs are taken from a list of genes shown to report phylogenetic distances between bacterial strains (Zeigler, 2003), with the additional constraint that they comprise a single copy in most of the 1108 genomes of our dataset.

### 1.3 Determination of the synteny network

The procedure for determining the set of gene pairs that are in a synteny property is explained in (Junier and Rivoire, 2013). Here we recall the main steps of this procedure.

Inter-gene distances – The distance between two genes is measured in base pairs, from the mid-point of their nucleotide sequence. To account for the fact that genomes may have several chromosomes, may be non-circular and have different lengths, we formally circularize linear chromosomes and normalize them to a common length *L* by setting all distances exceeding *L*/2 to *L*/2: if d is the actual distance in base pairs, we thus define a normalized distance *x* by *x* = min(1, 2*d*/*L*). The normalized distance between genes on distinct chromosomes is also set to *x* = 1. We take *L* = 500 kb, but our results are not sensitive to the exact value of this cutoff (the typical extension of the synteny segments that we find is far below 500 kb).

Genome weights – The number *M*_*ij*_(*x*) of genomes in which genes *i* and *j* are at normalized distance *x*_*ij*_ ≤ *x* is computed as *M*_*ij*_(*x*) = ∑_*g*_ *ω*_g_1(*x*_*ij*_ ≤ *x*), with genome weights defined by *ω*_*g*_ = 1/∣{*h* : *D*_*gh*_ < *δ*}∣, where ∣{*h* : *D*_*gh*_ < *δ*}∣ denotes the number of genomes *h* at phylogenetic distance at most *δ* from *g* (Morcos *et al*., 2011); this weighting procedure defines an effective number of genomes as *M*′ = ∑_*g*_ *ω*_*g*_.

Significance of proximity – Assuming a uniform distribution of genes along a circular genome of length *L*, the probability of observing a distance less than *xL*/2 between 2 given genes is just *x*. In this null model, the number *M*_*ij*_(*x*) of genomes with normalized distance *x*_*ij*_ ≤ *x* thus follows a binomial law B(*M*′, *x*), where *M*′ is the effective number of genomes. The probability *π*_*ij*_(*x*) of observing *M*_*ij*_(*x*) events is therefore *π*_*ij*_(*x*) = *I*_*x*_(*M*_*ij*_(*x*), *M*′ − *M*_*ij*_(*x*) + 1), where *I*_*x*_(*m*, *n*) is the regularized incomplete beta function. The least likely and therefore most significant normalized distance 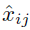 between a given pair of genes *ij*, is the one minimizing *π*_*ij*_(*x*), which defines 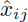 and an associated *p*-value 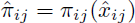.

Under the null model, the distribution of 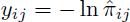 is found to have an exponential tail, *ψ*_0_(*y*) ~ *e*^−*ay*^, with an exponent *a* depending on *M*′ (Figure 13). Given a threshold of significance *π*\*, we compute the fraction *σ*_*s*_ of significant pairs, with 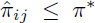, and estimate the fraction of false positive pairs as 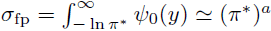. Imposing a false discovery rate FDR = *σ*_fp_/*σ*_s_ thus determines the threshold of significance *π*\*.

To treat pairs of COGs *ij* with multiple copies (genes), we fix a gene *g*_*i*_ in *i*, count the number *n* of genes in *j* at normalized distance less than *x* and compute the probability of the event as *p*(*x*) = 1 − (1 − *x*)^*n*^. The analysis is then performed as for *n* = 1 with 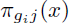 now standing for 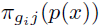, thus defining 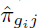. We then define 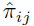 as the most significant observation when considering successively each gene *g*_*i*_ in *i*, i.e., 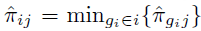. Different numbers of genes in *i* and *j* may imply 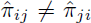. Pairs of proximal COGs are identified by requiring that both 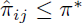 and 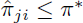.

### 1.4 Segments and long-range co-regulation

How much of gene co-expression between segments may be explained by “seg-regulation”? First, a significant fraction (∼ 25%) of highly correlated pairs of genes have a common seg-TF (Figure 9A), but only a small percentage have exactly the same set of seg-TFs (inset). Given the hierarchical organization of the regulatory network of *E. coli* (Ma *et al*., 2004), this is not surprising: having the same set of seg-TFs is a strong requirement, unlikely to be necessary for a form of seg-regulation to be present. Nevertheless, by retaining only the segments below a certain size, which ensures that the phenomena are not associated with the length of the segments, we loose the significant tendency for highly correlated genes to have a seg-TF (blue points in Figure 9C). Altogether, this shows that seg-regulation can drive co-expression (Figure 4D), but that this phenomenon play, overall, only a secondary role in regulating gene expression (Figure 9A). Incidentally, it also shows that the most correlated genes tend to belong to the largest segments, another noticeable feature.

Inter-segment co-expression must then rely on mechanisms that do not involve the action of TFs, at least not as regulators interacting with the RNA polymerase at the level of promoters (Browning and Busby, 2004). Several possibilities may be contemplated. For instance, we find that co-expressed genes tend to belong to segments with heEPODs (Vora *et al*., 2009) (Figure 9C). Yet, as for seg-TFs, the fraction of gene pairs with this property is low (∼ 10%); moreover, the tendency disappears when considering smaller segments. On the other hand, we observe that co-expressed genes are more likely to belong to segments that contain at least one region that is bound by Fis, a NAP that is involved in the supercoiling-mediated control of transcription initiation (Travers and Muskhelishvili, 2005) (Figure 9B); in this case, the tendency is more pronounced for small (< 4 kb) segments (Figure 9B). Altogether, this shows that seg-TF regulation is sufficient but not necessary for causing high co-expression, whereas “seg-Fis regulation” is necessary but not sufficient (∼ 50 % of the uncorrelated pairs of segments contain Fis binding regions).

